# Joint development recovery on resumption of embryonic movement following paralysis

**DOI:** 10.1101/2021.01.08.425893

**Authors:** Rebecca A. Rolfe, David Scanlon O’Callaghan, Paula Murphy

## Abstract

Fetal activity *in utero* is a normal part of pregnancy and reduced or absent movement can lead to long-term skeletal defects such as Fetal Akinesia Deformation Sequence (FADS), joint dysplasia and arthrogryposis. A variety of animal models with decreased or absent embryonic movements show a consistent set of developmental defects providing insight into the aetiology of congenital skeletal abnormalities. At developing joints defects include reduced joint interzones with frequent fusion of cartilaginous skeletal rudiments across the joint. At the spine defects include shortening and a spectrum of curvature deformations. An important question, with relevance to possible therapeutic interventions for human conditions, is the capacity for recovery with resumption of movement following short term immobilisation. Here we use the well-established chick model to compare the effects of sustained immobilisation from embryonic day (E) 4-10 to two different recovery scenarios: (i) natural recovery from E6 until E10 and (ii) the addition of hyperactive movement stimulation during the recovery period. We demonstrate partial recovery of movement and partial recovery of joint development under both recovery conditions, but no improvement in spine defects. The joints examined (elbow, hip and knee) showed better recovery in hindlimb than forelimb, with hyperactive mobility leading to greater recovery in the knee and hip. The hip joint showed the best recovery with improved rudiment separation, tissue organisation and commencement of cavitation. This work demonstrates that movement post paralysis can partially-recover specific aspects of joint development which could inform therapeutic approaches to ameliorate the effects of human fetal immobility.

**Summary Statement:** The study reveals that embryonic movement post paralysis can partially-recover specific aspects of joint development, which could inform therapeutic approaches to ameliorate the effects of restricted fetal movement *in utero*.

## Introduction

Reduced Fetal Movement (RFM) is a common clinical presentation in obstetric practice, with 22-25% of women perceiving decreased fetal movement resulting in poor perinatal outcomes (reviewed in Lai et al., 2016; Dutton et al., 2012). RFM *in utero* is associated with a number of conditions and syndromes including Fetal Akinesia Deformation Sequence (FADS) which represents a spectrum of defects in bone and joint formation including hypomineralised, brittle bones prone to fracture (Temporary Brittle Bone Disease), and contracture of joints (reviewed in Shea et al., 2015); joint dysplasia, particularly of the hip (reviewed in Nowlan, 2015); and arthrogryposis, defined as multiple joint contractures, affecting approximately 1 in 3000 live births (Skaria et al., 2019; Hall, 2014). Effects of RFM are variable and can range from mild to severe depending on the developmental window in which movement is interrupted (Filges et al., 2019). Short term absence of fetal movements at approximately 8 weeks of gestation, lasting over 3 weeks, has been theorised to be sufficient to result in the clinical features of arthrogryposis (Kowalczyk and Felus, 2016). The multiple contractures in arm and leg joints that result are associated with an increase in connective tissue around the immobilised joints, curvature abnormalities of the spine including kyphosis and scoliosis and disuse wastage of the muscles that mobilise joints (Ma and Yu, 2017; Hall, 2014). In most cases the reasons behind reduced fetal movement are unknown but the use of patient specific case studies of rare movement disorders (e.g. Prader-Willi syndrome), in combination with retrospective studies, further highlight the causative relationship between diminished fetal movements and skeletal anomalies (Donker et al., 2009; Bigi et al., 2008; Fong and De Vries, 2003; Moessinger, 1983).

The use of animal models has allowed direct investigation of the impact of reduced movement on skeletogenesis and has established that mechanical forces produced by embryonic movements are crucial for normal skeletal development (reviewed in (Rolfe et al., 2018; Felsenthal and Zelzer, 2017; Shea et al., 2015; Nowlan et al., 2010b)). Animal immobilisation models include pharmacological paralysis of muscle (in chick and zebrafish models), and genetic lesions that result in muscle absence or immobile muscle (in mouse and zebrafish models). Immobility results in specific effects on synovial joints, including reduction of the interzone region between adjacent skeletal rudiments, with continuity of cartilaginous rudiments across joints (fusion) in many cases; loss of normal cellular organisation with absence of the chondrogenous layers at the ends of rudiments (zones of future articular cartilage marked by increased cell density oriented parallel to the joint line); and failure to commence cavitation (Singh et al., 2018; Nowlan et al., 2014; Roddy et al., 2011b; Nowlan et al., 2010a;Kahn et al., 2009; Osborne et al., 2002). Changes within the rudiment termini also result in abnormal joint shape (Sotiriou et al., 2019; Brunt et al., 2016; Brunt et al., 2015; Roddy et al., 2011b) and all of these changes have been shown to be underpinned by altered gene expression and activation of signalling pathways that guide essential developmental steps including Wnt, BMP and Hippo (Shea et al., 2019; Rolfe et al., 2018; Singh et al., 2018; Brunt et al., 2017; Rolfe et al., 2014; Roddy et al., 2011b; Kahn et al., 2009). Disturbances of the spine due to immobility include curvature abnormalities, posterior and anterior vertebral fusions and altered vertebral shape (Levillain et al., 2019; Rolfe et al., 2017; Hosseini and Hogg, 1991). In clinical conditions and experimental animal models, the timing of initiation and duration of immobilisation is critical for the phenotypic abnormalities that result.

The development of both the axial and appendicular skeletons is sensitive to immobilisation in animal models from very early stages, from as early as embryonic day (E) 3 in the chick (Bridglal et al., 2020; Rolfe et al., 2017; Roddy et al., 2011b). A number of studies have monitored movement of the chick embryo through developmental time reporting amniotic and embryonic movements from as early as E3, however independent limb movements were not reported to take place until E5 or E6 (Wu et al., 2001; Oppenheim, 1975; Hamburger and Balaban, 1963). Given the clear effects of immobilisation on limb bone and joint development at time points prior to reported movement, here we address this apparent conundrum by looking specifically at the possibility of limb movements between E4 and E6.

While we know that short term embryonic immobility results in skeletal abnormalities, little is known about the capacity for the system to recover if movement resumes following short term immobilisation. There are indications that some aspects of the system can, at least partially, recover. Infants with Temporary Brittle Bone Disease (TBBD) can recover bone strength by normal mechanical stimuli in the first year of life. Joint shape abnormalities in infants are shown to be somewhat plastic, for example if congenital developmental dysplasia of the hip is identified early, joint shape can be ‘reset; by harnesses (reviewed in Vaquero-Picado et al., 2019). A recent study used physical external manipulation of hip joints in immobilised chick embryos, and showed more normal joint morphogenesis compared to unmanipulated contralateral limbs (Bridglal et al., 2020). This important question has implications for the long-term potential for recovery in conditions caused by fetal immobilisation and the potential development of therapies, either *in utero* or post-natal, to ameliorate the effects of restricted movement.

Here we use the chick model to investigate the effects of resumption of movement post paralysis and the potential to recover from skeletal abnormalities caused by short term immobilisation in a variety of limb joints and the spine. We compare two potential recovery scenarios; 1) where embryos are left to recover naturally following short term rigid paralysis through administration of the widely used neuro-muscular blocking agent 0.5% Decamethonium Bromide (DMB) and 2) where paralysis is followed by treatment with 0.2% 4-aminopyridine (4-AP), known to cause hyperactivity and increases fetal movement (Pollard et al., 2016; Pitsillides, 2006). We assess movement in the embryo following the recovery period under both scenarios. While it is difficult to separate the effect of the short term duration of the immobilisation from potential amelioration due to a recovery period, the comparison of natural recovery and stimulation of hyperactive movement with 4-AP provides the opportunity to investigate response to different levels of resumed movement. We show that embryonic mobility partially resumes following a period of short-term immobilisation, both naturally and following hyperactive drug treatment, and, while partial recovery from immobilisation abnormalities is achieved in limb joints it is not achieved in the spine. Within limb joints there is greater recovery in the hindlimb than forelimb, especially following hyperactive movement induction. Findings from this study suggest that movement stimulation can ameliorate the effects of paralysis on joint development.

## Results

### Limb displacement occurs from stage HH23 (E4)

Given that immobilisation from E3-6 has strong effects on limb joint development, while limb movement has been reported to commence only at E6 (Wu et al., 2001; Hamburger and Balaban, 1963) we further examined embryo movement specifically between E3 and E6. We utilised video recording and frame-by-frame image analysis to assess movement events and record any limb displacement, precisely staging each embryo (Table 1). No embryo movement was recorded in any specimens observed at E3. The first body movements were recorded at E4, precisely at stage HH22 when 8/11 embryos observed showed bending of the embryo trunk, most usually in the sagittal plane, with a steady increase in movement events over subsequent stages until all embryos were motile within the 2-minute video timeframe by HH24. Limb movement was assessed by relative displacement of the limb, comparing across video stills (Table 1, column 5). Outline drawings across a movement event (1-2 seconds apart) were overlaid at the dorsal aorta and aortic arch to reveal the relative displacement of the forelimb.Clear and distinct limb displacement relative to surrounding landmarks (aorta, eye and dorsal surface of the embryo) was recorded from as early as HH23 (8/11 cases) and in all specimens from HH24. It is unclear if such movements are solely passive, caused by the bending of the trunk, or have any contribution from spontaneous contraction of forming limb muscle masses at the latter stages (myotubes are first detected at HH25 (Kardon, 1998)). From HH27 (at E6) limb movements become larger and more obviously independent, corresponding to earlier observations (Wu et al., 2001; Hamburger and Balaban, 1963).

**Table 1:** The onset of embryo movement during chick development from E4-E6; precise HH stages noted. Analysis of two-minute video recordings of each specimen, recording body movement and limb displacement. Column 5 shows example analysis (2 examples for each stage) where the dorsal aorta and aortic arch (a/aa) (black arrowhead) were overlaid and the limb outlined in successive still images 1-2 seconds apart; Green = initial position, blue= moved, orange = final position. Fl; forelimb, 1mm scale bar for each.

| Stage | No of embryos | Embryonic day | Movement | Overlay image analysis of forelimb displacement (2 independent examples/stage) |
| --- | --- | --- | --- | --- |
| HH17-19 | 6 | E4 | No movement |  |
| HH20 | 7 | E4 | No movement |  |
| HH21 | 4 | E4 | No movement |  |
| HH22 | 11 | E4<br>(10 E4, 1 E5) | 8/11 some body movement |  |
| HH23 | 11 | E4<br>(8 E4, 3 E5) | 9/11 body movement;<br>8/11 limb displacement |  |
| HH24 | 4 | E5<br>(3 E5, 1 E4) | 4/4 body movement<br>4/4 limb displacement |  |
| HH25 | 15 | E5<br>(14 E5, 1 E6) | 15/15 body movement<br>15/ 15 limb displacement |  |
| HH26 | 5 | E5/E6<br>(2 E5, 3 E6) | 5/5 body movement<br>5/5 limb displacement |  |
| HH27-29 | 16 | E6 | 16/16 body movement<br>16/16 large limb movements |  |

### Embryonic movement partially resumes following a period of short-term immobilisation, both naturally and following hyperactivity drug treatment

The effects of immobilisation by rigid paralysis using the neuromuscular blocking agent DMB on skeletal development have been previously documented across various treatment periods including detailed analysis of the effects on knee joint development with treatment between E4.5 and E7 (Roddy et al., 2011b), on hip development over a series of treatment times and durations with early immobilisation from E4 for 3 days being most disruptive (Bridglal et al., 2020; Nowlan et al., 2014) and effects on spinal development with treatments from E3 to E9 (Levillain et al., 2019; Rolfe et al., 2017). Here we examine the capacity for recovery from effects at multiple joints and the spine following early immobilisation between E4 and E6 analysed at E10. To definitively establish sensitivity to immobilisation across the time frame used in the recovery experiment, we first carried out a preliminary experiment where embryos received daily treatments either from E4-E6 (early), harvested at E7, or from E7-E9, harvested at E10 (Fig 1A). Both early and later treatment regimens resulted in abnormal joints compared with control embryos (mock treated) (Fig. 1B), as well as other typical effects previously described such as altered spinal curvature, rudiment length reduction and joint contracture (data not shown) (Rolfe et al., 2017; Nowlan et al., 2008). Whereas control treated specimens, staged as HH30 (E7) show clear separation of cartilaginous rudiments (Fig. 1Ba and d), immobilisation from E4 to E6, assessed at E7 (stage verified as HH30) show dramatic reduction in rudiment separation at both knee (Fig 1Bb and c) and hip joints (Fig 1Bd and f). Following later (E7 to E9) immobilisation, assessed at E10/ staged to HH36, there is also a clear reduction in the joint interzone and the separation of rudiments across the joint (Fig.1 Bh-i, Bk-l) and additionally, while signs of the commencement of cavitation are evident in control specimens at HH36, there is no sign of cavitation commencing in either knee or hip joints following immobilisation in specimens at the same stage (Fig. 1 Bg-l; yellow arrow in controls). This preliminary study established that even short, early immobilisation from E4 to E6 results in disturbance of development similar to more sustained immobilisation, as previously described.

**Figure 1:**
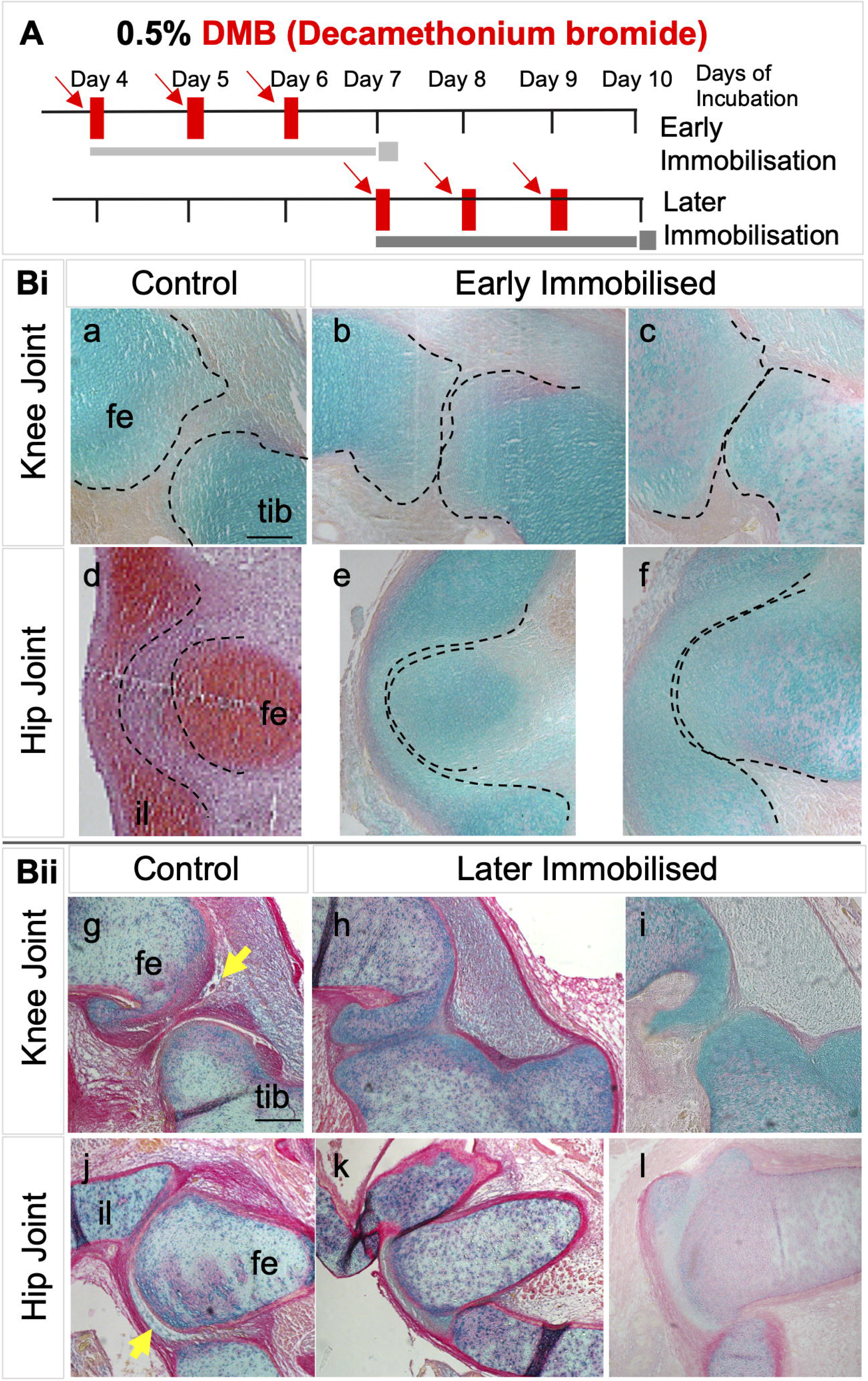
Both early and late immobilisation of chick embryos *in ovo* result in abnormal development of knee and hip joints. (A) Schematic of chick embryo immobilisation regimens using daily dosing for three consecutive days with 0.5% Decamethonium Bromide (DMB) as indicated by red arrows. Early; from day 4 to day 6; and later from day 7 to day 9. Specimens were harvested at E (embryonic day)7 (HH30), or E10 (HH36), respectively (each specimen staged). (B) Histological sections of knee and hip joints from early (i) and later (ii) immobilisation regimes as indicated. Representative replicate immobilised specimens are shown compared to control (b and c, e and f, h and i, k and j show replicate specimens). a-c, e-l are stained with alcian blue; d stained with Safranin-O. Dotted lines overlaid on the HH30 images outline the cartilage rudiments showing altered rudiment separation with immobilisation. Yellow arrows in HH36 indicate the initiation of cavitation in control knee and hip joints, absent with immobilisation. fe; femur, tib; tibiotarsus, il; ilium. Scale bar 100µm.

To assess if movement resumes following an initial period of DMB administration (0.5% DMB from E4 to E6), followed by either a period of natural recovery or hyperactivity stimulation (0.2% 4-aminopyridine (4-AP) daily, E7-E9) (Fig. 2A), each embryo from each of the treatment groups was observed and movement recorded over a 60 second period on E10, prior to harvest (Fig. 2B). The four-point classification system established here to score extent of movement *in ovo*, found that all embryos in the control group (n= 23) showed extensive movement with all but one embryo scored as having large body and limb bending movements at E10 (Fig. 2B). 64% of embryos subjected to sustained immobilisation from E4 to E9 (n= 14) (i.e. no recovery period) showed no movement (score of 0), two embryos had a score of one (minor body sway) and two a score of two (additional small limb movements), with only one embryo showing more extensive movement.

**Figure 2:**
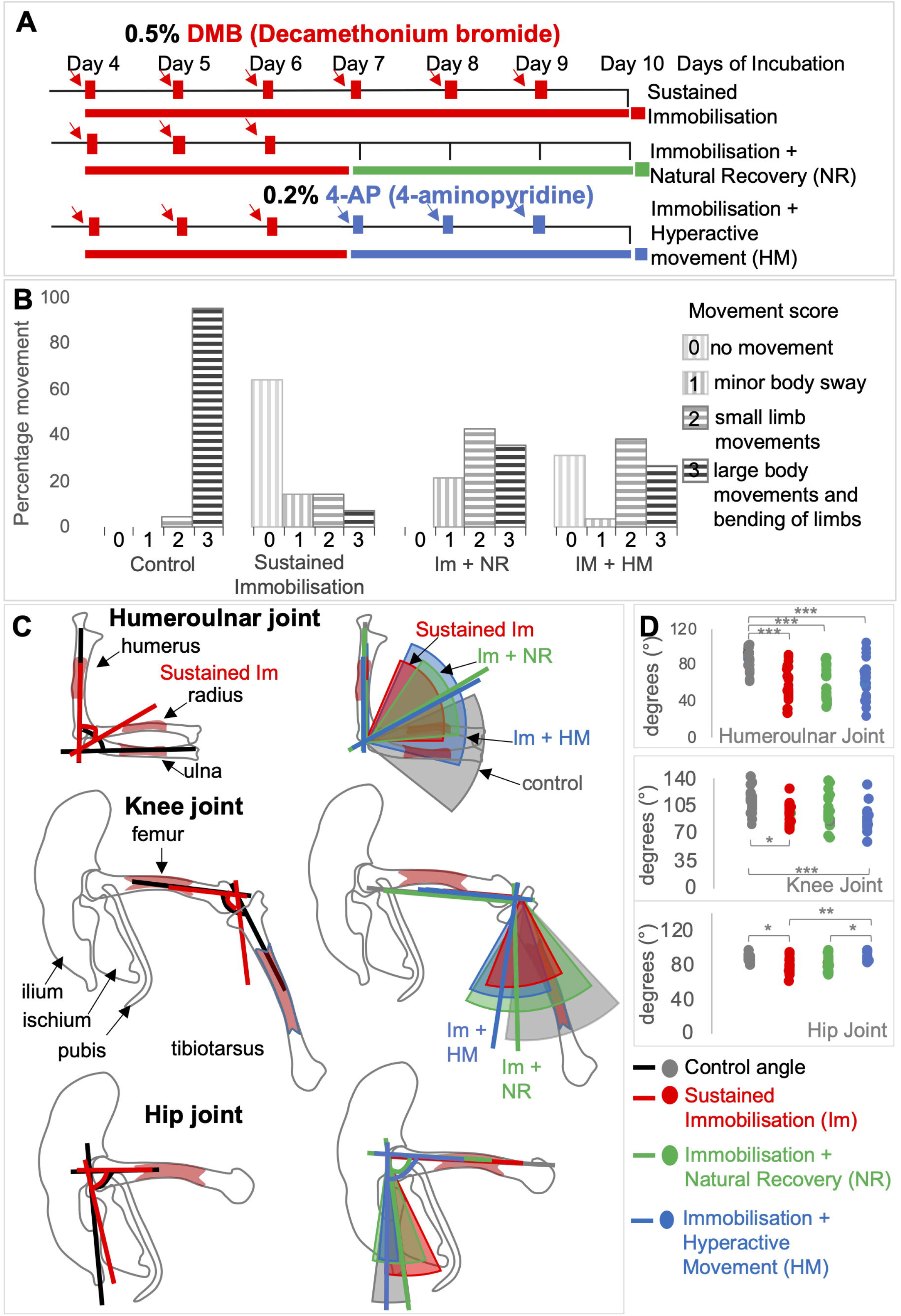
Embryonic movement resumes when early immobilisation (E4-E6) is followed by a natural recovery period (E7-E10; Im+NR) or induction of hyperactivity (4AP treatment; E7-E10; Im+HM). (A) Schematic of chick embryo immobilisation regimens using daily dosing with 0.5% Decamethonium Bromide (DMB) as indicated by red arrows commencing at day 4 of incubation; harvesting was at E10 under all regimens. Sustained immobilisation (red), treatment for 6 consecutive days; Immobilisation + NR (natural recovery) (green), treatment E4-E6; Immobilisation + HM (hyperactive movement treatment) (blue), treatment E4-E6 followed by addition of 0.2% 4-Aminopyridine (4-AP; blue arrows) on day 7 for 3 consecutive days. (B) Movement scores as indicated following observation of each embryo at E10 for 1 min periods (n=14-26 per group). Percentage of movements with scores 0-3 observed in each treatment group are shown. (C) Visual representation using schematic outline drawings for forelimb and hindlimb rudiments and joints, as labelled, with coloured lines indicating the mean joint angle for each group and the coloured segment overlay showing the angle range observed for each immobilisation regimen (Table 2). (D) Dot plots of joint angle including statistical analyses indicating the individual angles observed for each joint across groups. **Black** lines, **Grey** slices and dots; control ‘normal’ movement, **Red** lines, slices and dots; Sustained immobilisation (E4-E10), **Green** lines, slices and dots; Immobilisation (E4-E6) + natural recovery (E7-E10), **Blue** lines, slices and dots; Immobilisation (E4-E6) + hyperactive movement (E7-E10).*;p≤0.05,**;p≤0.01, ***;p≤0.001.

Both recovery groups showed increased movement in the post-immobilisation period compared to sustained immobilisation (p<0.01), with the majority in both groups having the highest movement scores of two or three, but were significantly less active than control embryos (p<0.05). All embryos in the immobilisation followed by natural recovery group (Im+ NR) showed some movement (79% scoring 2 (small limb movements) or 3 (large body and limb bending movements) (Fig. 2B) (n=14)). The recovery group where 0.2% aminopyridine was administered to stimulate movement displayed the greatest range of movement classifications and again with the majority scoring in categories two or three (Fig. 2B) (n=25). Overall, movement was significantly recovered following short periods of immobilisation, in both recovery groups, compared to sustained immobilisation although the extent of resumed movement was significantly less than in control embryos. There were no significant differences in movement scores between natural recovery or hyperactive stimulation (p=0.289) assessed in this way.

**Table 2:** Mean joint angles (+/-SEM) observed at the elbow, knee and hip in each group at E10. Numbers of replicates (n) indicated. Represented graphically and significance levels indicated in Fig. 2C and D.

| Joint | Control | Sustained<br>Immobilisation | Im + NR | Im + HM |
| --- | --- | --- | --- | --- |
| Humeroulnar | 88.8° ±3° n=26 | 59.7° ±4.1° n=20 | 59.7° ±3.6° n=23 | 62.6° ±4.9° n=20 |
| Knee | 110.2° ±3.2° n=23 | 93.7° ±3° n=18 | 97.2° ±4.7° n=19 | 87° ±4.4° n=15 |
| Hip | 86° ±1.2° n=15 | 77.7° ±2.3° n=15 | 80° ±2.1° n=18 | 87.9° ±1.6° n=10 |

Joint contractures are a common feature of rigid paralysis induced by DMB treatment so we assessed joint angle at elbow, knee and hip joints across the groups as an indirect indication of recovery from rigid paralysis following short term immobilisation. Comparing sustained immobilisation for 6 days to control showed abnormal flexion of all joints (Fig. 2C (red lines compared to black lines) as expected. Elbow joint angle for both recovery groups and sustained immobilisation was significantly more flexed than controls, (p≤0.001) (Fig. 2C-D, Table 2). In all immobilisation groups there is a large range of elbow joint angles observed (Fig. 2C-D) with the largest range for the immobilisation plus hyperactive movement group totalling in excess of 80 degrees (Fig. 2C (blue segment) and D (blue dots)). This group also showed the greatest variation in extent and types of movements observed (Fig. 2B). For the knee joint, sustained immobilisation also resulted in more flexed knee joints (p=0.012, Fig. 2C-D, Table 2) while knee joint angles following natural recovery (Im +NR) were not significantly different from the control group (p=0.066) (Fig. 2C green line in knee joint). The immobilisation with hyperactivity treatment group (Im + HM) on the other hand, were significantly flexed compared to controls (p<0.001, Fig. 2C-D), again with a large range in the data for recovery groups. Analysis of the hip joint (femur and ilium (posterior)) showed less variance in all groups and only hip angles under sustained immobilisation were significantly more acute than control joints (Fig. 2C-D). Hip joint angles were most similar to control when short-term immobilisation was followed by hyperactivity treatment (Fig. 2C-D, Table 2).

In summary, joint angle analysis corroborates the movement scores in indicating partial recovery of normal joint position following short term immobilisation. The effect was variable across joints with the greatest effect on restoration of normal hip joint angles under both recovery regimens; natural recovery resulting in more normal knee joint angles and the elbow joint remained most abnormally flexed and most similar to the situation under sustained immobilisation under both recovery scenarios.

### Effect of paralysis and recovery on skeletal development

#### i) Joint development

Abnormalities in chick knee and hip joint development following rigid paralysis have been well documented (Nowlan et al., 2014; Roddy et al., 2011b) while abnormalities at the elbow joint have also been noted (Roddy et al., 2011b) but not previously characterised in detail. Here we examine if a post paralysis recovery period can reduce or recover abnormalities observed under sustained immobilisation at the elbow, hip and knee. All groups were assessed at E10 (all verified at stage HH36), when early signs of cavitation are normally evident (Roddy et al., 2009). Using histological analysis of full series of sections through the joints of replicate specimens in each treatment category we assessed the elbow, knee and hip joint for evidence of recovery in three specific abnormalities caused by immobilisation; 1) reduced separation of the rudiments (reduced interzone) with partial fusion of cartilaginous rudiments at the joint in most cases (scored as presence or absence of joint fusion); 2) absence of distinguishable chondrogenous cell layers at the rudiment termini (altered tissue patterning) and 3) lack of initiation of cavitation indicated by the absence of a tissue free region within the joint.

Table 3 summarises the data across all joints while Figure 3 presents representative histological sections. Since the elbow joint consists of two sites of articulation, with the radius and ulna distal to the humerus, analysis of both the humeroradial (HRD) and humeroulnar (HUL) joints was performed separately (Table 3). All control joints at this stage displayed clear separation of the rudiments (Fig. 3, left hand column, red brackets) and characteristic tissue organisation at the joint interface and interzone; in particular the typical organisation of the chondrogenous layers (site of future articular cartilage) at the rudiment termini, evident as areas of increased cell density with orientation of cells parallel to the joint interface (Fig. 3, left hand column; yellow brackets). Early signs of cavitation are clear in all control joints as localised regions of tissue clearance (Fig. 3Aii, Bii, Cii, black arrows).

**Table 3:**
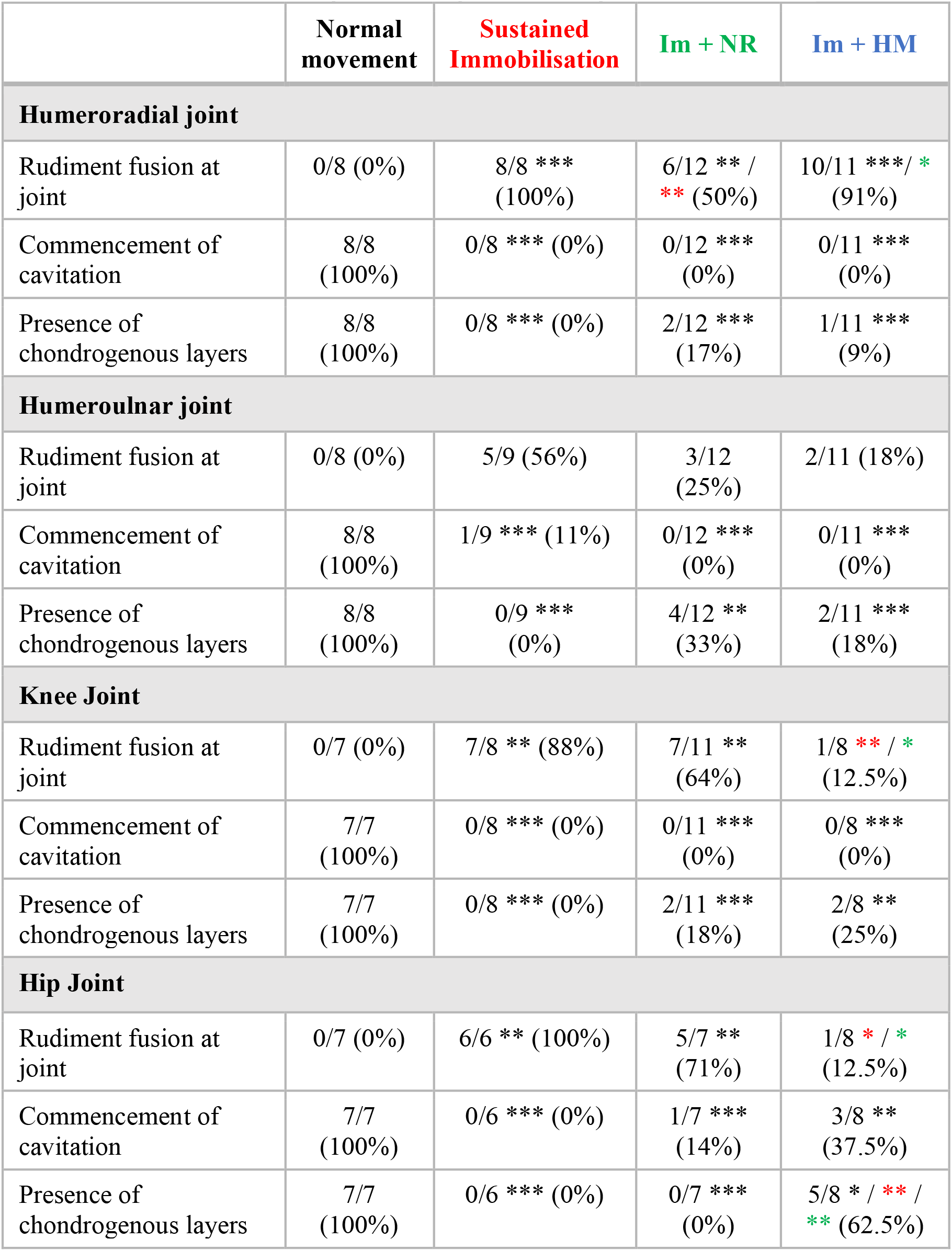
No. of specimens showing features of immobilisation at the elbow, knee and hip joints in control specimens (normal movement), following sustained immobilisation, and following recovery periods post-immobilisation; Natural Recovery (Im + NR) and Hyperactive Movement (Im + HM), as indicated. Colour of asterisks indicates the group comparison to which the significance level assessment relates, Black * to normal movement, Red * to sustained, Green * to Im+NR. *; p≤0.05, **; p≤0.01, ***; p≤0.001.

**Figure 3:**
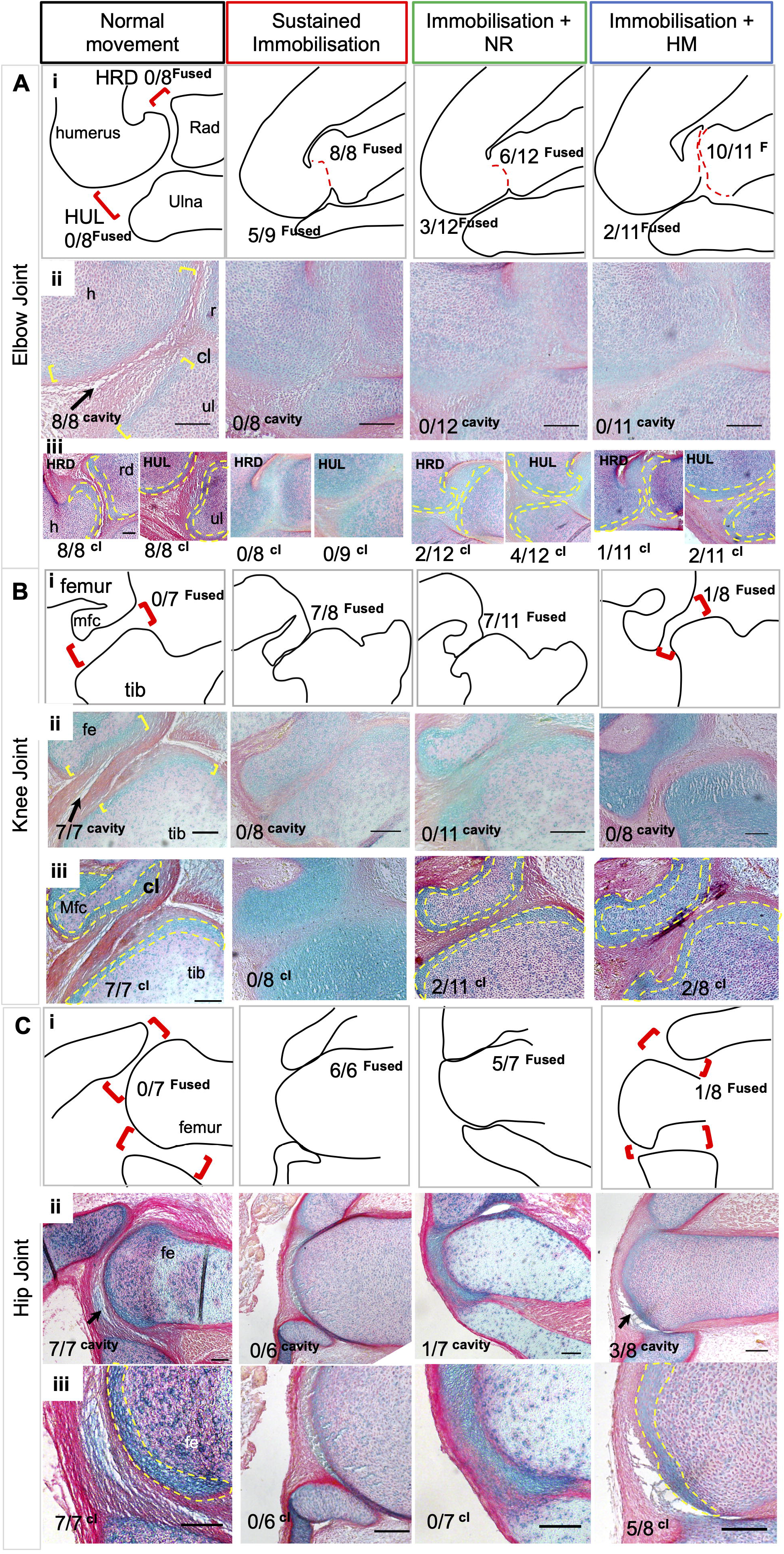
Elbow, knee and hip joint tissue patterning and morphogenesis are disrupted with sustained immobilisation, while movement resumption leads to partial recovery in aspects of joint organisation, as revealed by histological analysis. All joints were examined by longitudinal serial sections from medial to lateral of each specimen (n values indicated; representative images shown). Schematic outline drawings (row i) represent individual specimens from each experimental group as indicated (sections shown in row ii), through the elbow (A) knee (B) and Hip (C) joints. Row i: Red open brackets indicate normal rudiment separation/ interzone; red dotted lines indicate rudiment fusion (absence of interzone). The proportions of specimens with rudiment fusion observed across the groups is indicated. Row ii: histological sections as outlined in row i, indicating also the commencement of cavitation where visible (black arrow); numbers indicate number of specimens in each category where cavity commencement was observed. Row iii: chondrogenous layers, where present, are outlined in yellow dash; numbers indicate number of specimens in each category where chondrogenous layers (cl) are distinguishable. Abbreviations: HRD; Humeroradial, HUL; humeroulnar, h; humerus, rd; radius, ul; ulna, fe; femur, tib; tibiotarsus, Mfc; medial femoral condyle. All scale bars 1000µm.

#### Elbow joint

Following sustained immobilisation typical cellular organisation at the elbow joint is lost, similar to other limb joints, including separation between rudiments with rudiment fusion in 8/8 specimens analysed at the HRD interface, while fusion was observed in 56% (5/9) at the HUL (Fig. 3Ai, Table 3), suggesting a stronger effect of immobilisation on the HRD compared to the HUL at the elbow. This altered separation of the rudiments in immobilised joints is accompanied by absence of the clear organisation of cells within chondrogenous layers (0/8 and 0/9 at the HRD and HUL respectively under sustained immobilisation (Fig. 3Aiii). Complete absence of commencement of cavitation was also observed in both articulations of the elbow (0/8; Fig. 3Aii). With early immobilisation followed by a recovery period, rudiment fusion was evident at the elbow joint but in a lower proportion of specimens; 6/12 and 10/11 at the HRD; 3/12 and 2/11 at the HUL following normal recovery (Im +NR) and hyperactive movement (Im + HM) respectively (Table 3) with the humeroulnar joint again being impacted less (Fig. 3Ai, Table 3).

The presence of chondrogenous layers at rudiment termini also showed an indication of partial recovery with distinct chondrogenous layers in 2/12 HRD and 4/12 HUL joints with natural recovery (Fig. 3Aiii, yellow dotted lines indicating cellular territory), and 1/11 HRD and 2/11 HUL joints following recovery with hyperactive movement induction. However, neither recovery group reached a level of significant difference from sustained immobilisation in this respect. Despite improvement in rudiment separation and the presence of chondrogenous layers in some specimens following recovery periods however, there was no evidence of commencement of cavitation in either recovery group at this stage (Fig. 3Aii, Table 3).

#### Knee joint

Within the knee joint, there was 88% incidence of fusion between the medial femoral condyle and the tibiotarsus under sustained immobilisation and this was reduced with movement resumption; 64% (7/11) with natural movement and only 12.5% (1/8) with hyperactive movement where hyperactive movement recovery is statistically different to sustained immobilisation and is not different to the control situation (Fig. 3Bi, Table 3). In all joints immobilised for a sustained period there was complete absence of chondrogenous layers (0/8) and no evidence of commencement of cavitation (0/8) (Fig. 3Bii-iii). Resumption of movement resulted in the presence of chondrogenous layers in 4 specimens, 2/11 (18%) with natural recovery and 2/8 (25%) with induced hyperactive movement (Fig. Biii, Table 3). However, neither resumption of movement condition resulted in commencement of cavitation within this timeframe.

#### Hip joint

Analysis of the hip joint (articulation between the ilium and the femoral head) showed the best recovery with movement resumption. While again complete rudiment fusion, absence of chondrogenous layers and no evidence of commencement of cavitation were observed in all specimens with sustained immobilisation (Fig. 3C, Table 3), with movement resumption rudiment separation was observed in 29% of cases with natural movement and 87.5% with hyperactive movement (Fig. 3Ci, Table 3). While there was no apparent improvement in cellular organisation seen through the appearance of chondrogenous layers following natural movement resumption post paralysis, 62.5% (5/8) of cases showed recognisable chondrogenous layers following induction of hyperactivity post paralysis, highly statistically significant (Fig. 3Ciii, yellow dotted lines indicating region, Table 3). Unlike the other joints analysed, evidence of commencement of cavitation at this stage was observed in both movement resumption groups, 14% (1/7) with natural movement and 37.5% (3/8) with hyperactive movement (Fig. 3Cii, black arrow).

Taken together the data suggests that resumption of movement following short-term immobilisation can partly rescue the effects on joint development seen with sustained absence of movement, with greater recovery in the hindlimb than forelimb joints and generally greater recovery with hyperactive movement compared to natural resumption. Evidence of rudiment separation was observed in both hindlimb joints (knee and hip), with a greater incidence in the group stimulated for hyperactive movement post paralysis. The only immobilised joint to show evidence of commencement of cavitation at this stage following short-term immobilisation and a recovery period was the hip joint, while evidence of recovery of chondrogenous layer cellular organisation was seen in a proportion of specimens at all joints. At the elbow, the humeroulnar joint was slightly less impacted by sustained immobilisation and showed greater capacity for recovery than the humeroradial joint.

To corroborate the findings at limb joints, we examined another aspect of limb skeletogenesis previously shown to be sensitive to the loss of movement; skeletal rudiment length, measuring the length of the femur and the humerus across treatment groups (Fig. S1). While both rudiments showed significant reduction in length under sustained immobilisation, both again showed indications of partial recovery following resumption of movement. The femur showed no significant difference in length between control and immobilisation followed by hyperactive stimulation, while there was evidence of a trend with an increase in mean length in the humerus when immobilisation was followed by stimulation of hyperactivity, and while still significantly shorter than controls, a reduction in the significance level, compared to sustained immobilisation (Fig. S1, A, B, blue bars).

### ii) The spine

All groups of immobilised spines, including short-term immobilisation followed by a recovery period, with or without hyperactivity stimulation, were shorter than controls (p<0.001 in each comparison, Fig. 4A) with no differences in curved length between immobilised groups, sustained, natural recovery or hyperactive movement (Fig. 4A). Curvature deformities observed in the sagittal plane include hyperkyphosis and hyperlordosis, while abnormalities observed in the coronal plane are scoliosis (Fig. 5A schematic representations). Observed curvature abnormalities and associated reductions in spine length are seen in sagittal curvature outlines for each movement group (Fig. 4B).

**Figure 4:**
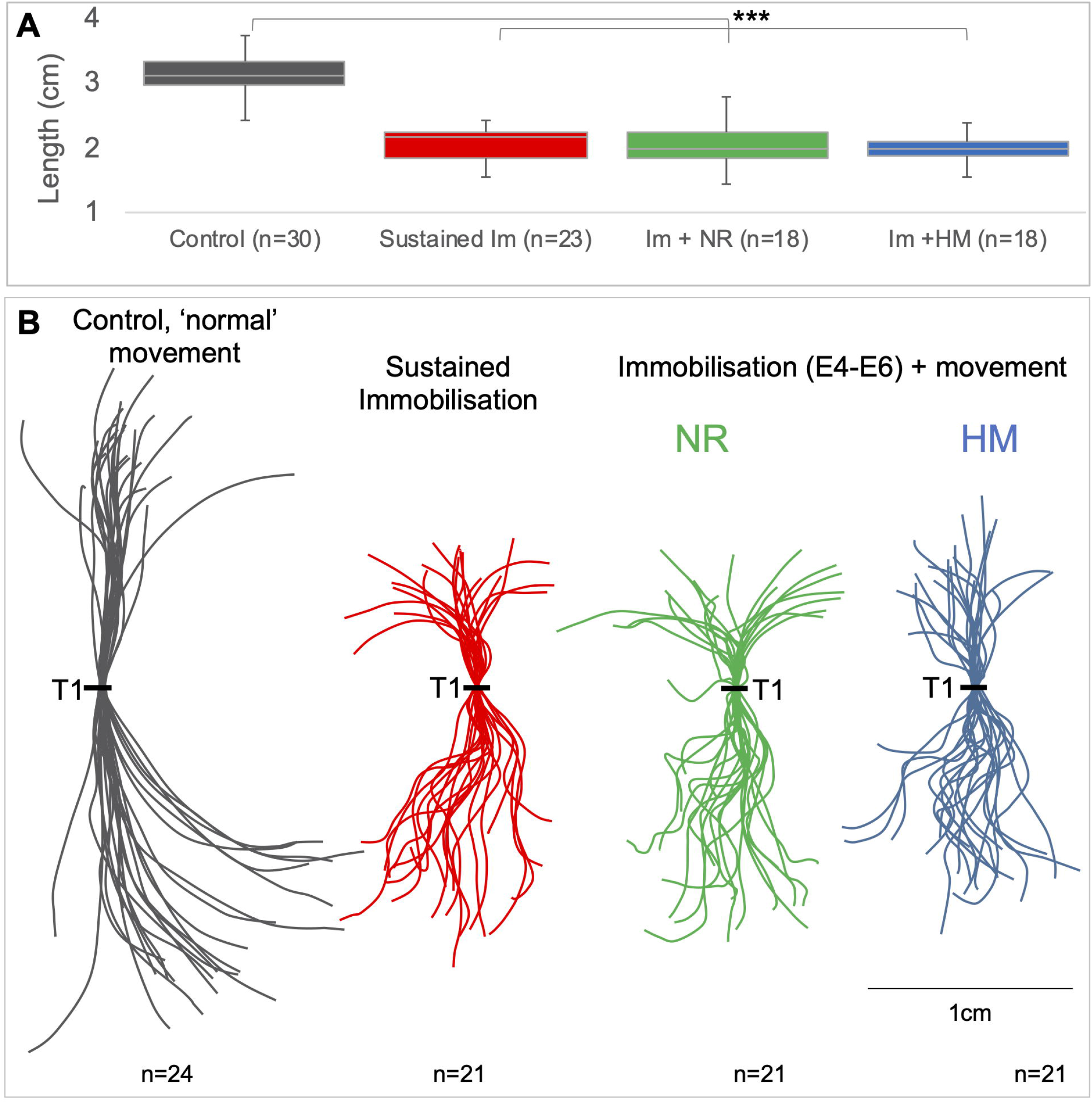
Spines from immobilised specimens with and without resumption of movement were shorter and abnormally curved compared to spines with normal movement. (A) Comparison of spine lengths in all movement groups ***p≤0.001, (B) Sagittal curvature outlines of control (grey), sustained immobilisation (red) and immobilisation followed by natural recovery (NR) of movement (green) or hyperactive movement (HM) (blue lines) show reduction in lengths and curvature abnormalities. Individual spines overlaid at thoracic vertebra 1 (T1). Scale bar 1cm. replicate numbers indicated.

**Figure 5:**
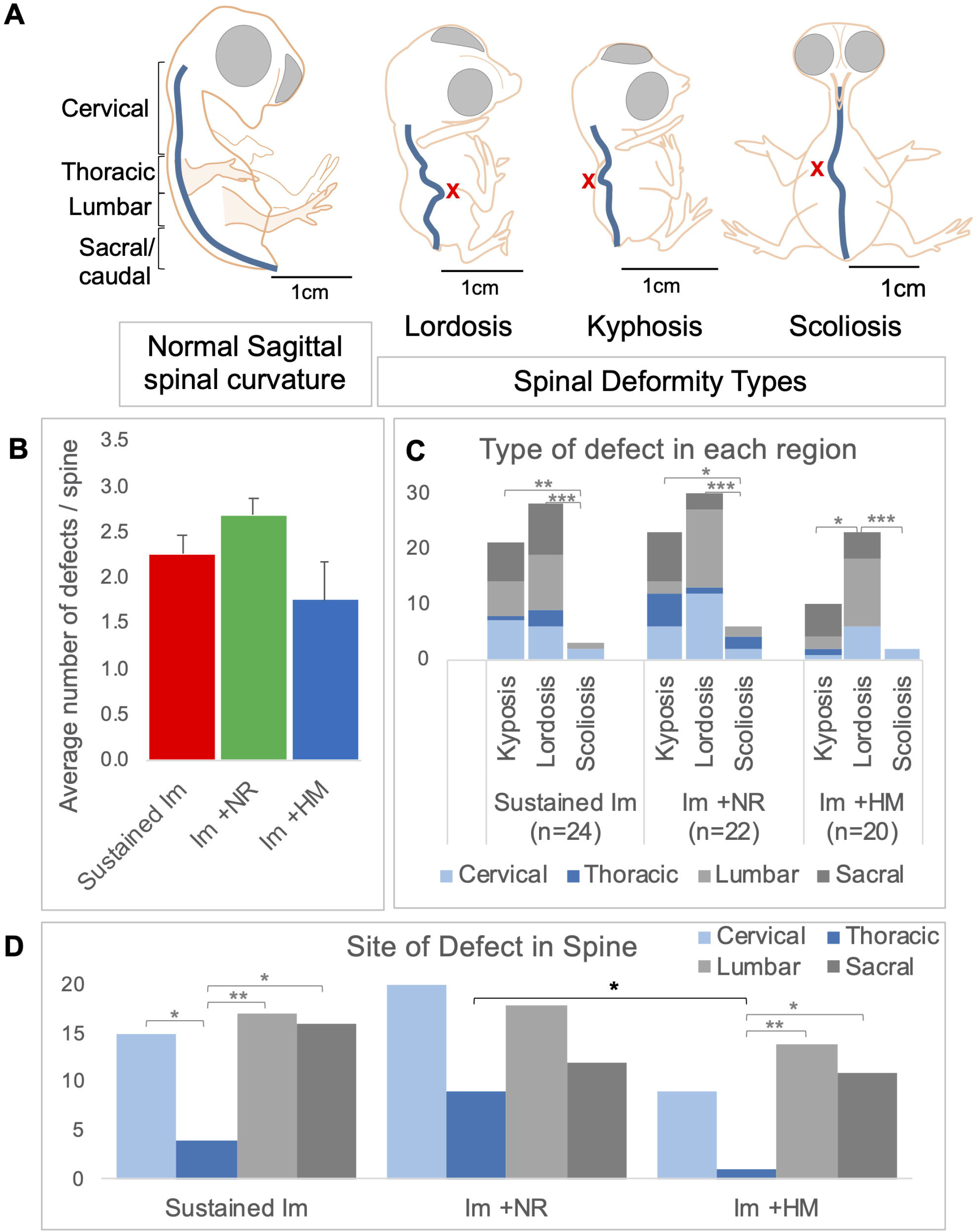
The cervical and lumbar regions are most highly affected by curvature deformities, hyperlordosis and hyperkyphosis, under immobilisation. (A) Schematic outlines of normal sagittal spinal curvature and the spinal deformities (X) of lordosis, kyphosis and scoliosis. (B) Average number of curvature defects observed in each reduced movement group. (C) Bar chart showing the number and type of spinal defects observed in each anatomical region (cervical, thoracic, lumbar and sacral) for all reduced movement groups. (D) Bar chart indicating the anatomical regions that are most affected by immobilisation *; p≤0.05,**; p≤0.01, ***; p≤0.001.

In all immobilisation regimens, there were 146 individual spinal deformities observed in 66 immobilised spines. Sustained immobilisation (E4-E10) for 6 days resulted in a total of 52 individual curvature deformities observed in 24 spines, with an average of 2.26 ± 0.22 (SEM) deformities per spine (Fig. 5B, red bar), while 59 individual defects in 22 spines were observed with natural recovery (NR) (an average of 2.68 ± 0.35 per spine, Fig. 5B, green bar) and 35 in 20 spines (an average of 1.75 ± 0.26) (Fig. 5B, blue bar) with hyperactive movement. There was no difference in the average number of defects per spine across immobilisation groups (Fig. 5B). Sustained immobilisation resulted in significantly more kyphotic and lordotic defects than scoliotic defects (p<0.001, Fig. 4C), totalling 21 incidences of hyperkyphosis, 28 incidences of hyperlordosis and 3 scoliotic bends, with natural recovery showing similar incidences and differences between defect type. Hyperactive movement resulted in significantly more lordotic defects than kyphotic and scoliotic (Fig. 4C). Combining the data, the most common abnormality was hyperlordosis at 55.5%, then hyperkyphosis at 37% and scoliosis was 7.5%, across all immobilisation regimens with and without a recovery period. Independent of immobilisation regimen, there were significantly more lordotic than kyphotic defects (p<0.045) or scoliotic (p<0.001) (Fig. 5C, data combined). The low incidence in scoliotic defects or abnormal curvatures in the coronal plane (11 incidences in 146) corresponds to previous observations following chick immobilisation (Rolfe et al 2017).

The cervical, lumbar and sacral anatomical regions were equally affected with regards to the total number of deformities observed with sustained immobilisation (15, 17 and 16 respectively), while the thoracic region was significantly less affected than other anatomical regions (Fig. 5D). The thoracic region was similarly significantly less affected than the lumbar and sacral regions with hyperactivity recovery (Fig. 5D) while no site-specific differences were recorded in the natural recovery group. Combining all immobilisation groups with or without a recovery period there were significantly more deformities in the lumbar and cervical anatomical regions, compared to the thoracic region, (p<0.004, p<0.016, respectively (2-way ANOVA)) (Fig. 5D).

Comparison of the recovery regimens reveals few differences except less defects in the thoracic region with hyperactive recovery of movement compared to natural recovery of movement (p=0.024 Fig. 5D, black bar).

## Discussion

This study advances our understanding of the plasticity of skeletal deformities caused by reduced embryonic movement by investigating the effects of resumption of movement post paralysis. It shows that movement resumption following rigid paralysis during early phases of skeletal development (from E4-E6) is only partially achieved, even with hyperactivity treatment, and this corresponds to partial recovery in some of the skeletal developmental defects caused by reduced movement. Recovery is seen in aspects of limb joint development but not in spinal defects. Within limb joints there is better recover in hindlimbs (hip and knee) compared to forelimbs (elbow), and overall better recovery with induction of hyperactive movement compared to natural resumption of movement. The hip joint showed the best recovery and closest to normal developmental progression post-paralysis. Overall, this demonstrates a degree of plasticity in terms of the dependency of normal joint development on embryonic movement and shows the potential for therapeutic intervention to improve outcomes in clinical joint abnormalities caused by reduced fetal movement such as in arthrogryposis and joint dysplasia.

Additionally, the study improves the chick immobilisation model as an experimental system to investigate the impact of reduced movement on development by refining knowledge on the commencement of movements in the embryo. Early studies by Hamburger and Balaban (1963) described commencement of body movement as early as E3.5 but they did not observe independent limb movements until E6.5. This 1963 study, often cited in the literature, creates a difficulty in understanding how immobilisation prior to E6 could have such strong impact on limb skeletal development, particularly on joint patterning (Fig 1). Subsequent studies have reported limb movement from E5 (Oppenheim, 1975) or E6 (Wu et al., 2001). We resolve this apparent conundrum by re-examining early embryo movements using video frame by frame analysis, combined with precise staging of embryos, establishing that distinct limb displacement occurs from HH23 (E4). While it is unclear if such movements are entirely passive, resulting from body movements propelled by spontaneous contraction of trunk muscles (Wu et al., 2001), or may have some contribution from spontaneous contraction of forming limb myotubes, detectable from HH25 (Kardon, 1998), they create a biophysical environment that could influence skeletal development, that would be altered in immobile specimens.

Skeletal abnormalities in infants caused by reduced fetal movement *in utero* are to some degree plastic, and therefore amenable to improvement by targeted therapeutics (reviewed in Vaquero-Picado et al., 2019; Miller and Hangartner, 1999). Directly investigating this important question of plasticity in animal immobilisation models is challenging, in particular the capacity for recovery following resumption of movement due to the difficulty of separating the effects of altered timing and duration of immobilisation and any recovery achieved following resumption of movement. To overcome this we compared two recovery scenarios, where embryos are allowed to recover naturally following short term rigid paralysis or where paralysis is followed by treatment with 0.2% 4-AP to increase fetal movement (Pollard et al., 2017). This level of 4-AP treatment is shown to result in an increase in frequency of movement by up to 175% (Pollard et al., 2016; Pitsillides, 2006) and to impact muscle structure, bone growth (Heywood et al., 2005) and tendon mechanical properties (Pan et al., 2018). Therefore, the effects of short-term immobilisation followed by a natural recovery period is not only compared to sustained immobilisation, but also to a recovery period with hyperactive movement. The finding that hyperactive mobility during the recovery period resulted in significantly greater recovery that the natural resumption of movement demonstrates that recovery is achieved, albeit partial. The achievement of recovery is also supported by the demonstration of the effects of the short-term period of immobilisation alone (E4-E6) which causes similar defects to sustained immobilisation with reduction of the knee and hip joint interzone (Fig 1). This same early period of immobilisation is also reported to have most severe effects on hip development (Bridglal et al., 2020). The recovery recorded here was variable across different aspects of skeletal defect analysed, as well as between natural resumption of movement and hyperactive movement, providing important insight into the potential and dynamics for recovery.

A further important aspect of the experimental design is the use of a simple movement scoring system to verify and assess movement in each of the experimental groups on the day of harvest, showing that movement does indeed resume following termination of immobilisation drug treatment, under both recovery regimens. Strikingly however movement does not return to levels seen in control embryos. One possibility explanation is that the developmental impact during the period of paralysis, including very severe alteration to tissue patterning, especially reduction in the interzone with partial cartilage fusion across the joint, may physically hinder free movement. Using this scoring system, movement following natural resumption and administration of the hyperactivity drug was not differentiated but this may be due to limited assessment; while movement event and type within a 60 second timeframe were recorded, movement duration or frequency was not assessed. Video recording and analysis, similar to that used here to record normal movements, would permit a more refined analysis but this would have delayed the harvesting and fixation of specimens, potentially affecting survival and compromising analysis and comparison of stage matched specimens. In combination with the movement classification scoring approach, we assessed joint angle as an indirect indication of movement resumption. Sustained immobilisation resulted in abnormal flexion of all joints as expected. There was a large range of angles recorded across the groups, especially following recovery with hyperactive movement, reducing the sensitivity of this approach for revealing significant differences between groups. However, this analysis corroborated the movement scores in revealing partial return to more normal joint position following short term immobilisation with the greatest effect on restoration of normal hip joint angles under both recovery regimens. In contrast the elbow joint remained most abnormally flexed under both recovery scenarios. This aligns well with the relative degree of recovery achieved in different joints.

Having established that embryonic motility partially resumes following short term immobilisation, we examined joint development, comparing the effects of both recovery scenarios to sustained absence of movement, and control specimens. Joint development is an important focus to assess the potential for recovery for two reasons: 1) the clinical relevance of joint developmental defects due to reduced fetal movement during pregnancy and 2) extensive characterisation of the effects of immobilisation on joint development in animal models, particularly the knee and hip joints in the chick (Bridglal et al., 2020; Sotiriou et al., 2019; Brunt et al., 2016; Nowlan et al., 2014; Roddy et al., 2011b; Nowlan et al., 2010a; Osborne et al., 2002). It was previously noted that chick elbow joints were affected similarly to knee joints (Roddy et al., 2011b) but here we describe elbow joint effects for the first time. To assess the potential for recovery upon resumption of movement we focussed on three aspects of joint developmental defects under immobilisation that could be readily scored on serial sections through the entire joint: 1) reduction in the joint interzone, scored as presence or absence of fusion between skeletal rudiments at the joint; 2) presence/absence of chondrogenous layers at rudiment termini; and 3) commencement of cavitation, denoted by tissue clearance. Using fusion between skeletal rudiments as a measure of the severity of effect showed some level of recovery following resumption of movement at all joints but most significantly at the hip and knee joints, particularly with hyperactivity induction post paralysis. It is important to note that absence of a fusion score indicates a less severe phenotype but does not necessarily indicate a normal interzone, where size might still be reduced. In previous studies 3D imaging was used to allow precise orientation, accommodating comparable measurements across immobilised and control specimens, showing reduction in the size of the interzone following immobilisation at the knee joint (Roddy et al., 2011b) and the hip (Bridglal et al., 2020). Here all specimens were analysed using serial histological sections so that all three aspects of joint development progression could be scored in each specimen. The difficulty of ensuring that the orientation of physical sections is the same across specimens makes comparable measurements impossible. The fusion score however gives a reliable indicator of recovery.

Chondrogenous layers form at rudiment termini, at the knee joint at HH32, clearly recognisable in histological sections by increased cell density with cell alignment parallel to the joint line (Roddy et al., 2009). They give rise to the articular cartilage of the future joint (Ito and Kida, 2000) and are molecularly distinct from the underlying transient cartilage that will be replaced by bone (Singh et al., 2018). Chondrogenous layers do not form in limb joints of both chick and mouse immobile embryos (Singh et al., 2018; Roddy et al., 2011b; Nowlan et al., 2010a; Kahn et al., 2009). Here we find that by far the best recovery in the appearance of chondrogenous layers is at the hip with hyperactivity (5/8 specimens compared to 0% at all joints examined under sustained immobilisation). Other joints, and all joints with natural resumption of movement, show limited recovery in small numbers of specimens. The third feature scored; initiation of cavitation at the joint, showed no indication of recovery in either the elbow or knee joint but is evident at the hip in three of eight specimens with hyperactivity treatment. Taking all three features together it is clear that the best recovery is seen at the hip joint with hyperactive movement, followed by the knee, with the elbow the least improved. It is interesting to note that the greatest improvement at the hip joint corresponds to resumption of a more natural angle in both recovery groups (Fig 2).

We have previously hypothesised that local biophysical stimuli generated from movement create a type of positional information that contributes to the correct patterning of emerging tissues in the joint (Roddy et al., 2011b). We have also shown changes in the molecular profiles and signalling pathways active across the territories of the joint (Shea et al., 2019; Rolfe et al., 2018; Singh et al., 2018; Rolfe et al., 2014). In particular, there is partitioning of signalling activity with BMP signalling active within the skeletal rudiment, at a distance from the joint interzone, and the canonical Wnt pathway active at the joint line, but this spatial restriction is lost in immobilised mouse and chick specimens (Rolfe et al., 2018; Singh et al., 2018). Cell territories are altered on multiple levels in immobilised specimens, also including localised cell proliferation patterns (Roddy et al., 2011b; Kahn et al., 2009; Germiller and Goldstein, 1997) and nuclear localisation patterns of YAP within skeletal rudiments, related to changes in shape at the joint interface (Shea et al., 2019). Cell migration is also an important feature of the forming joint (Shwartz et al., 2016) which may be another cellular activity affected by biophysical stimuli (Rolfe et al., 2018), of particular interest given the importance of cytoskeletal regulation during cell migration. The partial recovery of cellular organisation seen here indicates that the molecular mechanisms that control localised tissue differentiation, sensitive to biophysical stimuli generated by movement, can recover if appropriate biophysical stimuli resume, even partially. In this study we have not assessed molecular profile, cell proliferation or cell migration under recovery, with the focus here on profiling overall recovery according to reliable morphological markers, but this will be important to address in future studies.

Reduced movement clearly impacts multiple aspects of joint development with multiple molecular and cellular changes sensitive to embryo movement. We assess recovery across multiple facets of joint development progression which are interrelated but distinct. Whereas we separately assess cellular organisation within the joint territory (reduction of the interzone and appearance of chondrogenous layers) and commencement of cavitation, some other studies use the term cavitation to encompass these multiple aspects of joint development. Osborne et al. (2002) devised a cavitation score which encompassed a spectrum of effects from full rudiment fusion to cavitated joints. Bridglal et al. (2020) assess the effects of a range of immobilisation regimens on hip joint development using 3D image analysis which does not assess the different cellular aspects detailed here, referring to observed reduction in rudiment separation as a cavitation effect. An important aspect of the Bridglal et al. (2020) study is the use of manual manipulation to move one immobilised limb, elegantly showing clear improvement in rudiment separation at the hip of the manipulated limb compared to the contralateral immobilised limb. It is interesting that we see best recovery in the hip. While Bridglal et al propose that movement causes physical weakness at the joint leading to cavitation, we propose that biophysical stimuli affect multiple aspects of cellular behaviour, at molecular, cell shape and cell migration levels. Specifically focusing on cavitation, Dowthwaite et al. (1998, 2003) show that hyaluronan synthesis and distribution plays an important role in cavitation; molecular components of the system are altered in immobilised embryos (Roddy et al., 2011b; Dowthwaite et al., 2003).

Recovery is not achieved in the spine with no difference observed with and without resumption of movement: all immobilised spines, including the recovery groups, were shorter and abnormally curved with the most common defect observed being lordosis and the thoracic region the least affected overall. The defects observed here are in agreement with previous findings (Levillain et al., 2019; Rolfe et al., 2017) but extend the analysis by comparing deformity type, site and number following immobilisation, as well as examining the capacity for recovery. Hyper-kyphosis and hyper-lordosis were the most common defects observed with the cervical and lumbar regions the most affected. Congenital kyphosis can be caused by a failure of formation, or more commonly, a failure of segmentation of vertebrae, while congenital lordosis is caused by failure of posterior segmentation, or spinous process fusion (Lonstein, 1999). Posterior and anterior fusion of vertebrae was observed at curvature abnormalities in all immobilised groups, along the length of the spine (Data not shown), similar to earlier findings (Rolfe et al., 2017). The reduced impact on the thoracic region may be related to a stabilising effect of the ribs which have been shown to be independent of effects on thoracic vertebral shape or curvature associated with immobilisation (Levillain et al., 2019). The variability in recovery observed here, between the spine and the joints and indeed between different joints, brings important insight on the capacity for recovery and warrants further investigation to understand site-specific recovery better. The stark difference in recovery between spine and limb joints may be related to developmental timing differences. While formation of the sclerotome, from which the vertebrae emerge, begins at approximately E2.5 with early cartilage cell differentiation by E5 and distinct segmented cartilaginous vertebrae by E6 (Scaal, 2016; Scaal and Christ, 2004; Shapiro, 1992), limb skeletogenesis occurs relatively later (Pacifici et al., 2006). The critical period of short-term immobilisation in this study (E4-E6) therefore corresponds to relatively later events in the spine including the appearance of distinct cartilaginous vertebrae. Relative timing might also explain why there is better recovery seen in the hip joint compared to more distal joints, particularly with respect to commencement of cavitation. Since cavitation is the latest of the features scored to appear during normal development, and since there is a proximo-distal gradient in developmental timing along the limb, it is possible that examination at a later stage would show better recovery in the more distal elbow and knee joints. Another important consideration in understanding the variability in recovery is the level and type of normal movement involved. While biometric studies *in utero* have profiled curvature changes of the developing human spine (Choufani et al., 2009) analysis of *in ovo* spinal movements has not been performed. Also while embryonic limb movement has been captured and modelled (Verbruggen et al., 2018a; Verbruggen et al., 2018b; Verbruggen et al., 2016; Nowlan et al., 2012; Roddy et al., 2011a; Roddy et al., 2011b; Nowlan et al., 2008), no such studies to date have modelled axial movements in order to understand their role in spine development. One contributing factor to superior recovery observed at the hip joint upon resumption of movement may be the impact of both limb and body movements at the hip whereas distal limb joints are only impacted by isolated limb bending movements. Quantifying and separating the contribution of mechanical input from these sources would be of value to determine the contributory role they play in hip joint development. Incorporating technological advancements in movement analysis (Pollard et al., 2016) and alignment of individual embryo movements with recovery could further elucidate site-specific capacity for recovery.

The work presented here provides a detailed morphological description of response within the skeletal system to restoration of movement following a period of immobility. It is the first study to integrate analysis of the appendicular and axial skeleton providing insight into the differential plasticity of the skeletal system and potential for recovery. In particular it shows that multiple aspects of joint patterning, disturbed when mechanical stimulation is removed, can recover when movement resumes. Information from this research could inform clinical assessment of congenital conditions in which short periods of paralysis occur *in utero*.

## Materials and Methods

### Egg incubation and *In ovo* movement manipulation

Fertilised eggs (Ross 308, supplied by Allenwood Broiler Breeders), were incubated at 37.7°C in a humidified incubator. Work on chick embryos does not require a licence from the Irish Ministry of Health under European Legislation (Directive 2010/63/EU), all work on chick embryos was approved by the Trinity Ethics committee. Following 3 days of incubation, 5mls of albumen was removed from each egg using an 18-gauge needle. Immobilisation (rigid paralysis) treatments consisted of daily application of 100μl 0.5% Decamethonium bromide (DMB) (Sigma-Aldrich) in sterile Hank’s Buffered Saline (HBSS) (Gibco) plus 1% antibiotic/ antimycotic (aa) (Penicillin, streptomycin, amphotericin B; Sigma-Aldrich), dripped directly onto the vasculature of the chorioallantoic membrane through the “windowed” egg.

Sustained immobilisation with daily treatments from E4-E9, harvested at E10, was compared to post-paralysis recovery groups as follows: 1) Immobilisation (E4-E6) followed by natural recovery (E7-E10) designated Im + NR (natural recovery); 2) Immobilisation (E4-E6) followed by daily treatment with 0.2% 4-aminopyridine (4-AP; flurochem) in sterile HBSS plus aa, designated Im + HM (hyperactive movement), as represented in Figure 2. The experiment was repeated independently three times with between 3-14 replicate specimens per group per experiment. Early and late treatment groups consisted of daily immobilisation from E4-E6, harvested at E7, or daily immobilisation at E7-E9, harvested at E10 respectively (Fig 1). Controls were treated with 100ul of sterile HBSS plus aa.

Harvesting was performed by cutting the vasculature surrounding the embryo and placing it in ice cold Phosphate Buffered Saline (PBS). Each embryo was staged using Hamburger and Hamilton criteria (Hamburger and Hamilton, 1992). Spines and limbs were dissected and either fixed in 4% paraformaldehyde (PFA) in PBS at 4°C, dehydrated through a graded series of ethanol (ETOH)/ PBS (25%, 50% 75%, 1x 10min washes, followed by 2x 10 min washes in 100% ETOH) for wax embedding, or fixed in 95% ETOH for 48-72hrs for wholemount staining.

### Assessment of normal embryo movement during development

Fertilised eggs were windowed at day 3 of incubation for *in ovo* observation (n=9) or transferred into culture for *ex ovo* observation (n=74), as previously described (Rolfe et al., 2018; Schomann et al., 2013), on days ranging from 3 to 6. The *ex-ovo* situation provides for better viewing and video recording of embryo movement while the *in ovo* samples allowed for comparison, with no differences noted between *ex-ovo* and *in-ovo* movement observations. Embryos were video recorded daily from E3-E6 using an 8-megapixel camera. The camera was placed in a fixed position above each embryo and videos of two minute duration were captured. Embryos were staged using morphological criteria according to Hamburger and Hamilton (Hamburger and Hamilton, 1951). The occurrence and types of movement observed in each video were recorded. Consecutive frame-by-frame stills of each movement were analysed using ImageJ software. Limb displacement was revealed by changes in forelimb position relative to landmarks on the head and trunk (eye, dorsal margin, dorsal aorta). During each observed limb movement, three still images 1-2 seconds apart, prior to, during and following a movement were overlaid, aligning at the dorsal aorta and aortic arch to capture the extent of limb displacement. Similar analysis of the hindlimb was hampered by less consistent visibility but similar movements of the hindlimb were evident.

### Movement scoring in embryos following a period of immobilisation and recovery

Embryos were observed daily and movement or absence of movement noted during the treatment regimens (E4-E9); as expected immobilised specimens showed drastically reduced movement. Movement scoring was carried out prior to harvest at E10 where each embryo was observed continually for 60 seconds and all movements recorded based on a simple classification scoring metric from 0 to 3; from a minimal value of 0= no body movements, 1= minor body sway, 2= some small limb movements and body sway, to the highest movement score of 3= large body movements and obvious bending of limbs. Replicate numbers for each group across experiments were as follows; control ‘normal’ movement n=23, sustained immobilisation n=14, immobilisation followed by natural movement n=14 and immobilisation followed by hyperactive movement n=25.

### Histological analysis

One forelimb and one hindlimb from each specimen were processed for paraffin wax sectioning while the contralateral limbs were processed for whole limb analysis. A full series of longitudinal sections (8µm) were prepared through each entire limb. Sections were dewaxed and rehydrated, stained for cartilage with 0.025% alcian blue in 3% acetic acid (1 hour) followed by 1% picro-sirus red (1 hour) for collagen, or 0.1% Safranin-O (1 hour). Individual, entire limb joints were assessed for 1) separation (or continuity) of cartilaginous rudiments at the joint (fusion); 2) the presence of chondrogenous layers (region of future articular cartilage) at rudiment termini at the joint interface i.e. organised cell layers typified by increased cell density with cells aligned parallel to the joint interface (Singh et al., 2016; Mitrovic, 1977); and 3) the commencement of cavitation indicated by the appearance of a tissue free region within the joint. A full series of sections, from medial to lateral, was evaluated for each joint. Longitudinal sections (8µm) of spines from each movement group were processed as above to assess vertebral separation.

### Whole skeletal preparation and imaging

Ethanol fixed whole limbs, and spines were stained for cartilage in 0.015% Alcian Blue in 95% Ethanol (in 20% glacial acetic acid) for 4–8 hours, followed by 0.01% Alizarin red in 1% Potassium Hydroxide (KOH) for bone and cleared in 1% Potassium Hydroxide (KOH) for 1– 6 hours. Whole spines and limbs were aligned for lateral view and photographed using an Olympus DP72 camera and CellSens software (v1.6). Measurements were made from 2-dimensional images using ImageJ. Qualitative analysis of spinal curvature and spinal deformities was performed, and quantitative assessment of spine and rudiment length and joint angle were measured.

### Spine height and deformity quantification

Spine length from cervical vertebra 1 (C1) to the last sacral vertebra was measured as a curved line through the centre of the vertebral bodies, from the most cranial to the most caudal, using the measurement function of ImageJ (v.1.51h).

To assess spinal curvature a line was traced along the centres of the vertebral bodies from the sagittal aspect to obtain an outline trace of sagittal curvature, as previously described (Rolfe et al., 2017). Sets of curvature outline traces were aligned at thoracic vertebra 1 (T1) and regions of pronounced kyphosis and lordosis were identified. Quantification of the number, type and sites of spinal deformities were assessed from whole stained spines from sagittal and coronal aspects. Replicate numbers for spine lengths and spinal deformities in each group were as follows; control ‘normal’ movement n=30, sustained immobilisation n=24, immobilisation followed by natural recovery n=22 and immobilisation followed by hyperactive movement n=20.

### Rudiment length and joint angle quantification

Cartilage and bone stained images of limbs were used to measure rudiment length and joint angle. For rudiment length replicate numbers across immobilisation and control groups were between 16 and 26. Quantification of joint angles was performed in both the forelimb and hindlimb; elbow (both the humeroulnar and humeroradial), knee and hip. All joints were observed from the lateral aspect and straight lines drawn through the longitudinal mid-point of the ossification site (observed with Alizarin Red). For example, in the knee joint a straight line overlay was drawn along the midline length of the femur and another straight line overlay on the tibiotarsus. The angle where the lines intersect (the vertex), was measured (Fig. S2.). Replicate measurements for each joint across all groups were as follows; humeroulnar n=20-26, humeroradial n=19-24, knee joint n=15-23, hip joint n=10-18 (range represents the different experimental groups).

### Statistical analysis

Statistical analysis was performed using SPSS (SPSS Statistics v26, IBM, corp.). To assess differences in movement scores, in mean joint angle, in spine lengths, in rudiment lengths, in joint defects, in the type and site of spinal deformities across and within experimental groups, univariate multiple comparisons analysis of variance (ANOVA) followed by Tukey *post-hoc* tests were used. To assess spinal deformities and sites of deformities with immobilisation a multivariate ANOVA followed by Tukey *post-hoc* tests were used. For all comparisons *p*≤0.05 was considered statistically significant.

## Acknowledgements

The authors are grateful to Dr. Claire Shea, Ms. Sarah Austin, Ms, Anna Shishparenok and Ms. Rachel Grant for their assistance with this research.

## Author contributions

Conception and design: R.A.R, P.M, Acquisition of Data: R.A.R., D.S.O’C. Analysis and interpretation of data: all authors. Writing the article: R.A.R, P.M. Final approval for publication: all authors.

